# NVX-CoV2373 ancestral and NVX-CoV2540 BA.5 protein nanoparticle vaccines protect against Omicron BA.5 variant in Syrian hamsters

**DOI:** 10.1101/2023.08.07.552330

**Authors:** Traci L. Bricker, Astha Joshi, Nadia Soudani, Suzanne M. Scheaffer, Nita Patel, Mimi Guebre-Xabier, Gale Smith, Michael S. Diamond, Adrianus C. M. Boon

**Affiliations:** Department of Medicine, Washington University School of Medicine in St. Louis, MO 63110, USA; Department of Pathology and Immunology, Washington University School of Medicine in St. Louis, MO 63110, USA; Novavax, Inc., Gaithersburg, MD 20878, USA; Department of Microbiology, Washington University School of Medicine in St. Louis, MO 63110, USA

**Author notes:** Corresponding author. Adrianus C. M. Boon.

## Abstract

The emergence of SARS-CoV-2 variants with greater transmissibility or immune evasion properties has jeopardized the existing vaccine and antibody-based countermeasures. Here, we evaluated the efficacy of boosting with the protein nanoparticle NVX-CoV2373 or NVX-CoV2540 vaccines containing ancestral or BA.5 S proteins, respectively, in mRNA-immunized pre-immune hamsters, against challenge with the Omicron BA.5 variant of SARS-CoV-2. Serum antibody binding and neutralization titers were quantified before challenge, and viral loads were measured 3 days after challenge. Compared to an mRNA vaccine boost, NVX-CoV2373 or NVX-CoV2540 induced higher serum antibody binding responses against ancestral Wuhan-1 or BA.5 spike proteins, and greater neutralization of Omicron BA.1 and BA.5 variants. One and three months after vaccine boosting, hamsters were challenged with the Omicron BA.5 variant. NVX-CoV2373 and NVX-CoV2540 boosted hamsters showed reduced viral infection in the nasal washes, nasal turbinates, and lungs compared to unvaccinated animals. Also, NVX-CoV2540 BA.5 boosted animals had fewer breakthrough infections than NVX-CoV2373 or mRNA-vaccinated hamsters. Thus, immunity induced by NVX-CoV2373 or NVX-CoV2540 boosting can protect against the Omicron BA.5 variant in the Syrian hamster model.

**IMPORTANCE:** As SARS-CoV-2 variants continue to emergence, the efficacy of prior and updated COVID-19 vaccines need to be tested. Here, we tested the efficacy of two nanoparticle protein-based vaccines in pre-immune hamsters against a challenge with the BA.5 Omicron variant of SARS-CoV-2. Compared to an mRNA vaccine boost, the nanoparticle vaccine NVX-CoV2373 and NVX-CoV2540 induced higher serum antibody binding and neutralization responses against ancestral Wuhan-1 or BA.5 variants. One and three months after the last immunization, hamsters were challenged with the Omicron BA.5 variant. NVX-CoV2373 and NVX-CoV2540 boosted hamsters showed reduced viral infection in the nasal washes, nasal turbinates, and lungs compared to unvaccinated animals. Animals that received the homologous vaccine, NVX-CoV2540, had fewer breakthrough infections than NVX-CoV2373 or mRNA-vaccinated hamsters. Together, our data shows that the BA.5 nanoparticle vaccine is effective and that it is important to update the COVID-19 vaccine to match currently circulating strains of SARS-CoV-2.

## INTRODUCTION

Severe acute respiratory syndrome coronavirus 2 (SARS-CoV-2) has caused hundreds of millions of infections worldwide and nearly 7 million deaths. Vaccines targeting the SARS-CoV-2 spike protein were developed within one year of the start of the pandemic. Several of these (mRNA, subunit, and adenoviral-vectored) are remarkably effective in protecting against severe coronavirus disease 2019 (COVID-19), with efficacy rates ranging from 75 to 95% depending on the vaccine, the circulating strain, and age of the individual ^1–3^. Vaccines also protect against infection antigenically closely related strains, albeit at lower (50-70%) rates ^4,5^.

In November of 2021, the Omicron variant of SARS-CoV-2 emerged, quickly spreading globally and replacing previous variants of concern (VOC) of SARS-CoV-2. Omicron variants harbor more than 30 amino acid substitutions in the spike (S) protein, which results in evasion of antibody immune responses and escape from protection of the original vaccines ^6^. In serum from individuals immunized with Ad26.COV2.S, mRNA-1273, or BNT162b2 vaccines, differences in neutralization comparing historical and pre-Omicron variants of SARS-CoV-2 with Omicron BA.1 or BA.5, ranged from 15-20 fold depending on the study population and vaccine ^7–11^. Although a third dose of ancestral S protein vaccines reduced this fold difference in neutralization, regulators in the United States and European Union approved bivalent vaccines containing both the original S and the BA.1 or BA.5 variant of the S protein. In humans, a boost with bivalent mRNA vaccines was 37% more effective in preventing severe disease and hospitalization than a monovalent booster ^12^. Bivalent SARS-CoV-2 mRNA vaccines increased the breadth of neutralization and protected mice against challenge with the BA.5 Omicron variant of SARS-CoV-2 ^13^. However, it is not clear if a bivalent or Omicron-specific vaccine is required for all COVID-19 vaccines including adenoviral-vectored and subunit-based nanoparticle vaccines.

Novavax developed a SARS-CoV-2 recombinant spike (S) protein nanoparticle vaccine comprised of the full-length prefusion trimers of S, and co-formulated with a saponin-based adjuvant, Matrix-M (NVX-CoV2373). In pre-clinical studies in mice and non-human primates, NVX-CoV2373 was effective against a homologous challenge with SARS-CoV-2 ^14,15^. Similarly, in mice, a version of the NVX vaccine based on the Beta (B.1.351) variant was effective against heterologous challenge with the Omicron BA.1 variant of SARS-CoV-2 ^16^. Currently, no pre-clinical studies have been published with Novavax’s BA.5-based vaccine (NVX-CoV2540). In humans, immunization with NVX-CoV2373 was effective against mild, moderate, or severe COVID-19 in clinical trials ^17–19^. Several trials reported the vaccine efficacy against symptomatic infection of 96% for the ancestral strain of SARS-CoV-2 and 86% for the alpha (B.1.1.7) variant ^19,20^. Boosting with a third or fourth dose of NVX-CoV2373 reduced the antigenic distance between the ancestral and Omicron BA.4/5 variants of SARS-CoV-2 ^21,22^, suggesting that repeated exposure to a subunit vaccine containing ancestral S protein induces a cross-reactive and cross-neutralizing antibody response.

Here, we evaluated the immunogenicity and efficacy of two protein-based nanoparticle vaccines, NVX-CoV2373 with the ancestral S protein and NVX-CoV2540 with the BA.5 S protein of SARS-CoV-2, in Syrian hamsters previously immunized twice with an mRNA vaccine (BNT162b2). We show in hamsters that heterologous boosting with the NVX vaccines induced robust immunity against Omicron BA.1 and BA.5 variants that protected against challenge with the BA.5 variant.

## RESULTS

### Immunogenicity of NVX-CoV2373 or NVX-CoV2540 in Syrian hamsters

Groups of 5-6 week-old male Syrian hamsters (n = 65) were immunized intramuscularly twice at four-week intervals with 5 µg of freeze-thawed lipid-encapsulated mRNA vaccine (BNT162b2) encoding a proline-stabilized Wuhan-1 S protein (**Fig S1A**) or PBS, and serum was collected 21 days after the first (day 21) and second (day 49) dose of the vaccine. Approximately one month later, the mRNA immunized animals were split into three groups and boosted with 5 µg of BNT162b2 mRNA vaccine (mRNA), 1 µg of the nanoparticle protein-based NVX-CoV2373 vaccine, or 1 µg of NVX-CoV2540 vaccine (n = 14-15 per group). The NVX-CoV2373 and NVX-CoV2540 vaccines contain the S proteins of Wuhan-1 or BA.5 variant of SARS-CoV-2, respectively. Serum was collected 21 days later and S-specific antibody responses were quantified by ELISA. As expected, serum from age-matched control hamsters that received PBS did not bind to the S protein by ELISA (**Fig S1B**). In comparison, serum collected from hamsters 21, 49 and 105 days after immunization and boosting contained anti-Wuhan-1 and anti-BA.5 S antibodies (**Fig 1 and S1**). Twenty-one days after the second mRNA immunization (day 49), the GMTs were 1:15,505, 1:10,266, and 1:7,700 (**Fig 1A**) for Wuhan-1 S in the three groups. These increased to 1:68,763 (4.4-fold, *P* < 0.0001), 1:193,237 (18.8-fold, *P* < 0.0001), and 1:114,139 (14.8-fold, *P* < 0.0001) after boosting (third dose) with mRNA, NVX-CoV2373, or NVX-CoV2540, respectively. These same sera were also tested for anti-BA.5 S antibodies. The GMTs for BA.5 S were 1:14,825, 1:9,167, and 1:9,412 (**Fig 1A**) in the three groups after two mRNA doses, and these increased to 1:33,966 (2.3-fold, *P* < 0.001), 1:78,603 (8.6-fold, *P* < 0.0001), and 1:72,360 (7.7-fold, *P* < 0.0001) after boosting with mRNA, NVX-CoV2373, or NVX-CoV2540, respectively. After boosting, the GMTs against Wuhan-1 and BA.5 S were higher in the mRNA, NVX-CoV2373, and NVX-CoV2540 immunized animals than the PBS control animals (*P* < 0.0001 for Wuhan-1 and *P* < 0.0001 for BA.5, **Fig 1B**). No significant differences were observed for serum antibody binding to Wuhan-1 or BA.5 S between NVX-CoV2373 and NVX-CoV2540 boosted hamsters.

**Figure 1.**
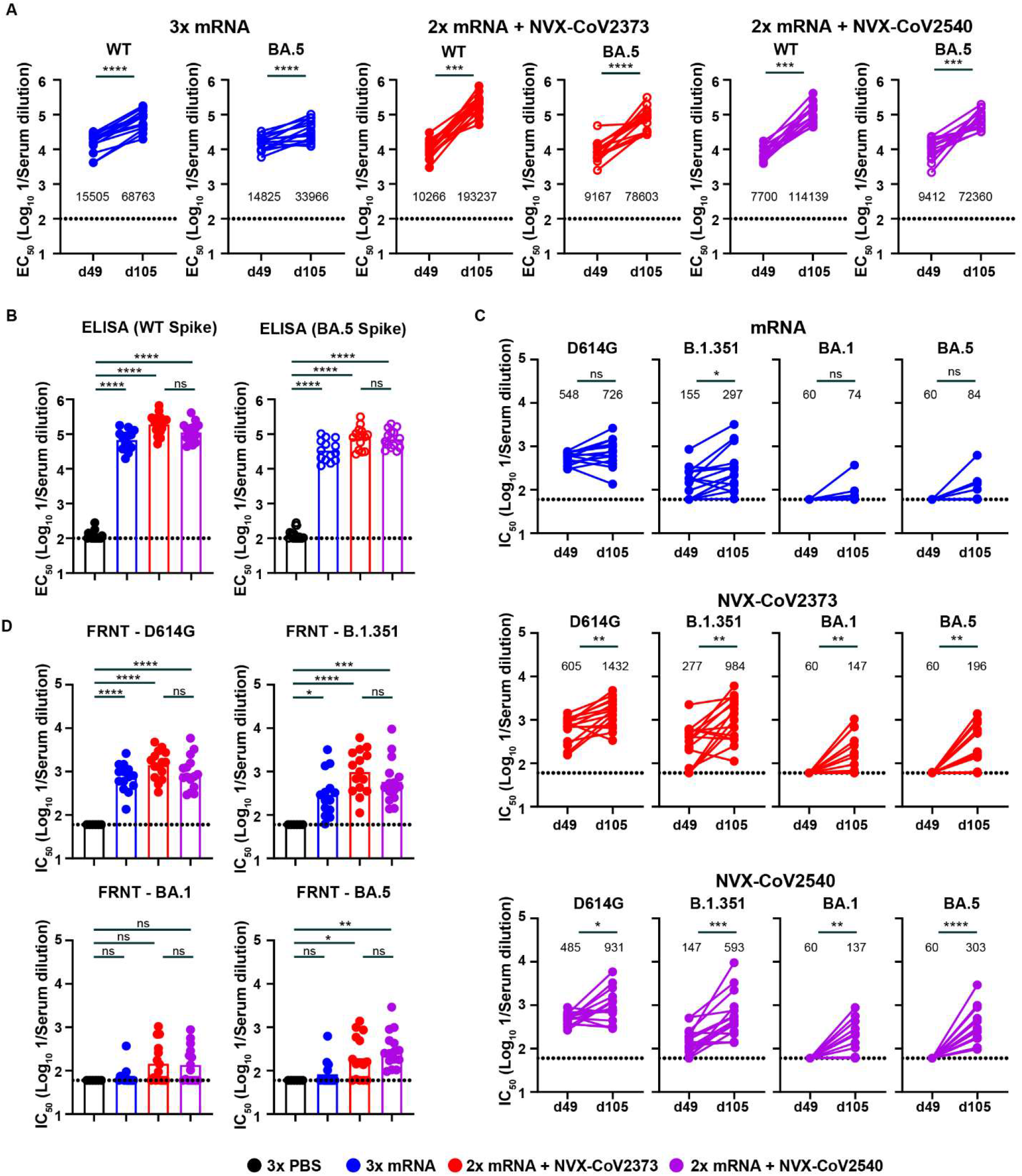
Boosting with NVX-CoV2373 and NVX-CoV2540 in previously mRNA-vaccinated hamsters induce high levels of serum antibodies. (**A**) Serum anti-Wuhan-1 and BA.5 S protein antibody response (EC_50_) in hamsters immunized twice with 5 µg of mRNA vaccine and boosted with mRNA (blue symbols), 1 µg of NVX-CoV2373 (red symbols), or 1 µg of NVX-CoV2540 (purple symbols). Serum was collected 21 days after the second dose or third dose of the vaccine. (**** *P* < 0.0001, *** *P* < 0.001 by paired two-tailed t-test). (**B**) Comparison of the serum antibody titers (EC_50_) against Wuhan-1 or BA.5 S protein measured by ELISA after a third immunization with mRNA (blue symbols), NVX-CoV2373 (red symbols), or NVX-CoV2540 (purple symbols). (**** *P* < 0.0001, *** *P* < 0.001, ** *P* < 0.01, ns = not significant by one-way ANOVA with a Šídák post-test on transformed EC_50_ values). (**C**) Serum neutralizing antibody responses (IC_50_) against WA1/2020-D614G, B.1.351, BA.1, or BA.5 SARS-CoV-2 in hamsters immunized twice with 5 µg of mRNA vaccine and boosted with mRNA (blue symbols), NVX-CoV2373 (red symbols), or NVX-CoV2540 (purple symbols) vaccines. Serum was collected 21 days after the second and third dose of the vaccine. (**** *P* < 0.0001, *** *P* < 0.001, ** *P* < 0.01, * *P* < 0.05, ns = not significant by paired two-tailed t-test on ln-transformed IC_50_ values). (**D**) Serum neutralizing antibody titer (IC_50_) against D614G, B.1.351, BA.1, and BA.5 variant of SARS-CoV-2, measured by FRNT, after a third immunization with mRNA (blue symbols), NVX-CoV2373 (red symbols), or NVX-CoV2540 (purple symbols). (**** *P* < 0.0001, *** *P* < 0.001, ** *P* < 0.01, * *P* < 0.05, ns = not significant by one-way ANOVA with a Šídák post-test on ln-transformed IC_50_ values). (**B and D**) Bars indicate the geometric mean values, and dotted lines are the limit of detection of the assays. Animals at the limit of detection are arbitrarily assigned this value. These values are combined with those having values above the limit to determine the GMT. Each symbol represents an individual animal. The results are from one experiment, and each symbol represents an individual animal (n = 14-15 for mRNA, NVX-CoV2373, and NVX-CoV2540, and n = 18-21 for PBS). See also **Figure S1**.

Serum samples collected 49 and 105 days after the first immunization (21 days after the second and third dose) were tested for neutralization of SARS-CoV-2 by focus reduction neutralization test (FRNT) against WA/1/2020-D614G (D614G), B.1.351, BA.1, and BA.5 strains of SARS-CoV-2 (**Fig 1C-D, and S1C**). Whereas serum from the PBS control animals did not neutralize SARS-CoV-2 (**Fig S1C**), serum from hamsters immunized with two or three doses of mRNA, or two doses of mRNA followed by one dose of 1 µg NVX-CoV2373 or NVX-CoV2540 did. The serum neutralization titers against D614G were 1:548, 1:605, and 1:485 in the mRNA, NVX-CoV2373 and NVX-CoV2540 groups prior to boosting, and these respectively increased to 1:726 (1.3-fold, *P* = 0.10), 1:1,432 (2.4-fold, *P* < 0.01), and 1:931 (1.9-fold, *P* < 0.05) after boosting (**Fig 1C-D**). No significant differences in GMTs of neutralization against the D614G virus was observed 21 days after boosting with NVX-CoV2373 and NVX-CoV2540 (**Fig 1D**). For B.1.351, the serum neutralization titer before boosting were 1:155, 1:277, and 1:147, and these increased to 1:297 (*P* < 0.05), 1:984 (*P* < 0.01), and 1:593 (*P* < 0.001) in the mRNA, NVX-CoV2373, and NVX-CoV2540 immunized hamsters respectively (**Fig 1C-D**). The difference in B.1.351-neutralizing GMTs was not statistically significant between the NVX-CoV2373 and NVX-CoV2540 boosted hamsters (**Fig 1D**). Two doses of mRNA did not induce detectable virus neutralizing antibodies against BA.1 or BA.5 (**Fig 1C-D**). Following boosting (third dose) with mRNA, or NVX-CoV2373 or NVX-CoV2540, the GMTs of neutralization increased to 1:74 (*P >* 0.05), 1:147 (*P* < 0.01), and 1:137 (*P* < 0.01) against BA.1, and 1:84 (*P* > 0.05), 1:196 (*P* < 0.01), and 1:303 (*P* < 0.0001) against BA.5 (**Fig 1D**). Boosting with NVX-CoV2540 induced neutralizing antibody titers in 100% of the animals, but the GMT was not significantly different compared to NVX-CoV2373 boosting (**Fig 1D**).

### Protection of NVX-CoV2373 and NVX-CoV2540 against BA.5 challenge in Syrian hamsters

Next, we challenged the boosted hamsters with 2.5 × 10^4^ plaque forming-units (PFU) of BA.5, a dose that enables robust replication in the upper and lower respiratory tracts. Animal weights were recorded daily for three days before nasal washes, nasal turbinates and lungs were collected for viral burden analysis (**Fig 2A**). No significant difference in weight was observed in the unvaccinated control (PBS) animals after BA.5 virus challenge (**Fig 2B**), which is consistent with the previously described attenuation of Omicron variants in rodents ^23^. Similarly, no differences in weight was observed in the immunized animals.

**Figure 2.**
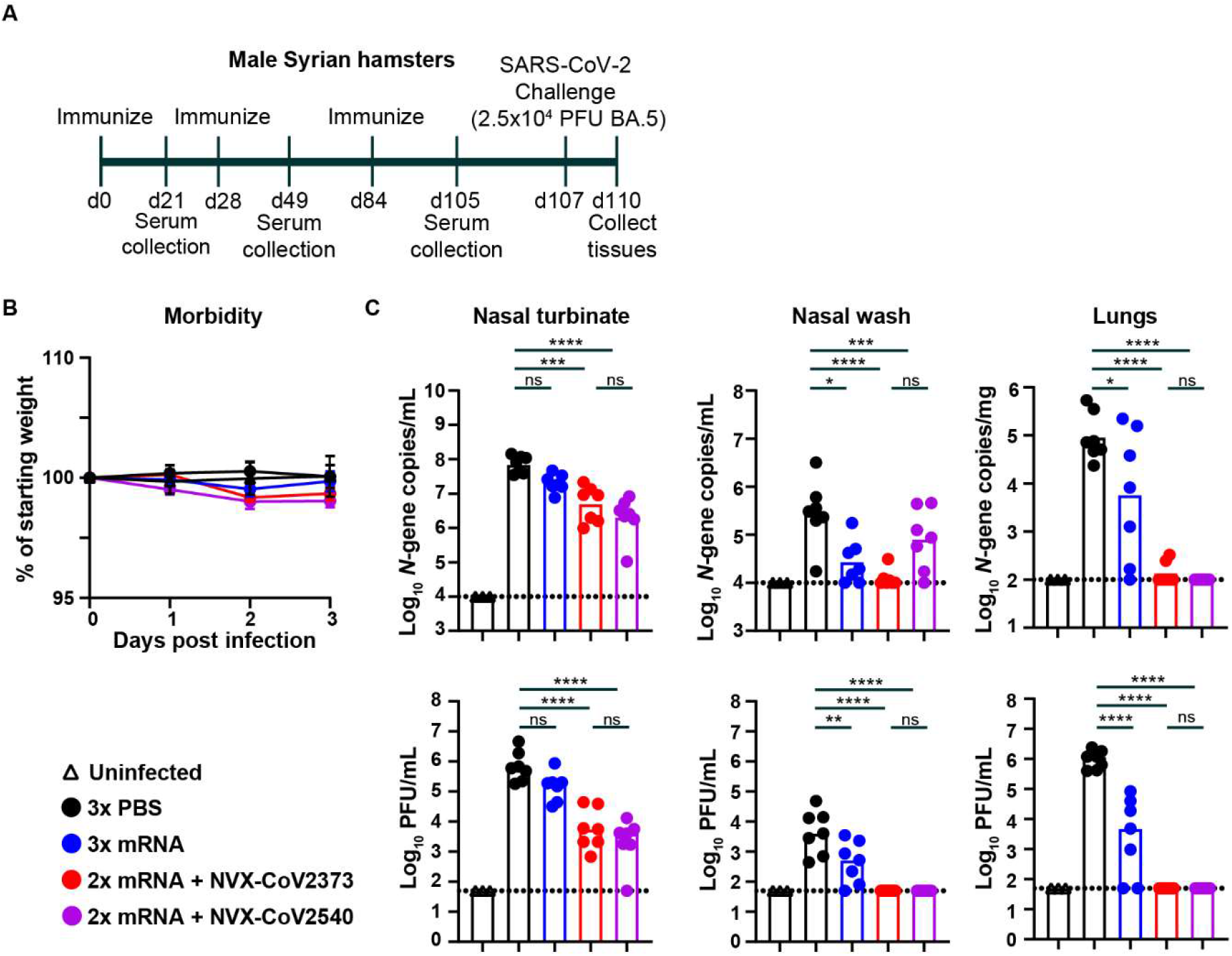
NVX-CoV2373 and NVX-CoV2540 boosters protect Syrian hamsters against challenge with BA.5. (**A**) Experimental design. (**B**) Mean ± SEM of weight loss/gain in SARS-CoV-2 challenged mice. (**C**) Syrian hamsters immunized twice with 5 µg of mRNA vaccine (BNT162b2) and boosted with 5 µg of mRNA (blue symbols), 1 µg of NVX-CoV2373 (red symbols), or 1 µg of NVX-CoV2540 (purple symbols), were challenged with 2.5 x 10^4^ PFU of BA.5, and nasal turbinates, nasal washes and lungs were collected for analysis of viral RNA levels (top panels) by RT-qPCR and infectious virus (bottom panels) by plaque assay (**** *P* < 0.0001, *** *P* < 0.001, ** *P* < 0.01, * *P* < 0.05, ns = not significant by one-way ANOVA with Šídák post-test on ln-transformed data). Uninfected control hamsters (open diamonds) were included as negative controls. Bars indicate the geometric mean values, and dotted lines are the limit of detection of the assays. Animals at the limit of detection are arbitrarily assigned this value. The results are from one experiment, and each symbol represents an individual animal (n = 7-8).

In terms of viral burden, no viral RNA or infectious virus was detected in our mock-infected control hamsters, as expected (**Fig 2C**). In unvaccinated control animals challenged with BA.5, we detected ∼10^8^ *N*-gene copies per mL of nasal turbinate homogenate, and this was reduced 3.4-fold (*P* = 0.12), 14-fold (*P* < 0.001), and 34-fold (*P* < 0.0001) in hamsters that received mRNA, NVX-CoV2373, or NVX-CoV2540 boosters, respectively. No significant difference was detected in viral RNA between NVX-CoV2373 and NVX-CoV2540 boosted animals. For infectious virus measurements, compared to control animals where we detected ∼7 × 10^5^ PFU/mL, an mRNA vaccine booster did not significantly reduce infectious virus titers (*P* = 0.20, **Fig 2C**). In contrast, boosting with NVX-CoV2373 and NVX-CoV2540 significantly reduced infectious virus levels in the nasal turbinate compared to the unvaccinated control group (120-fold, *P* < 0.0001 and 320-fold, *P* < 0.0001 for NVX-CoV2373 and NVX-CoV2540, respectively). No difference in infectious virus titers was observed between NVX-CoV2373 and NVX-CoV2540 boosted animals.

In the nasal washes of control groups challenged with the BA.5 virus, ∼3 × 10^5^ *N* gene transcript copies per mL and ∼4 × 10^3^ PFU/mL were measured (**Fig 2C**). Immunization with mRNA, NVX-CoV2373 or NVX-CoV2540 reduced *N* gene copy number 6.6-fold for mRNA (*P* < 0.05), 27-fold (*P* < 0.0001) for NVX-CoV2373, and 18-fold (*P* < 0.001) for NVX-CoV2540. Hamsters that received an mRNA vaccine booster had reduced *N* gene copy number (10-fold, *P* < 0.05, **Fig 2C**). Boosting with NVX vaccines also reduced infectious virus levels in the nasal washes below the limit of detection, whereas boosting with mRNA vaccine did not. No difference in infectious virus titers of viral RNA was observed between NVX-CoV2373 and NVX-CoV2540 boosted animals.

Viral burden also was measured in the lungs. In the control group, we detected ∼10^5^ copies of the *N* gene transcript per mg of lung tissue (**Fig 2C**). Boosting with mRNA vaccine resulted in reduced viral RNA levels by 16-fold (*P* < 0.05, **Fig 2C**), whereas boosting with NVX-CoV2373 (∼700-fold, *P* < 0.0001) or NVX-CoV2540 (925-fold, *P* < 0.0001) reduced the amount of *N* gene to near or below the limit of detection. Infectious virus titers were also measured. In unvaccinated control animals, we detected ∼10^6^ PFU/mL in the lungs, and this was reduced 370-fold (*P* < 0.0001, **Fig 2C**) in hamsters that received an mRNA vaccine booster. In contrast, in hamsters that received NVX-CoV2373 or NVX-CoV2540 as the booster, infectious virus levels were below the limit of detection (∼20,000-fold, *P* < 0.0001 for both groups).

### NVX-CoV2540 provides long-term protection against BA.5 challenge in Syrian hamsters

We next evaluated the long-term efficacy of the NVX vaccines against challenge with Omicron BA.5, as durability of vaccine responses have not been studied extensively in hamsters. Hamsters immunized with a booster (third) dose of mRNA, NVX-CoV2373 or NVX-CoV2540, were housed for another 13 weeks and a 4^th^ serum sample (day 174) was collected for serum antibody analysis (**Fig 3A**). The GMTs against Wuhan-1 S were reduced 5.7-fold (1:16,935), 5.1-fold (1:36,379), and 3.1-fold (1:36,650) between days 105 and 174 for mRNA, NVX-CoV2373, and NVX-CoV2540, respectively. The fold reduction in GMTs for BA.5 S was 3.2-fold (1:17453), 3.1-fold (1:21297), and 1.9-fold (1:39145) for these same sera (**Fig 3B-C**). The GMTs against Wuhan-1 and BA.5 S 13 weeks after mRNA, NVX-CoV2373, and NVX-CoV2540 boosting remained significantly (*P* < 0.0001) different compared to unvaccinated control animals. No difference in GMTs between NVX-CoV2373 and NVX-CoV2540 boosted animals was observed.

**Figure 3.**
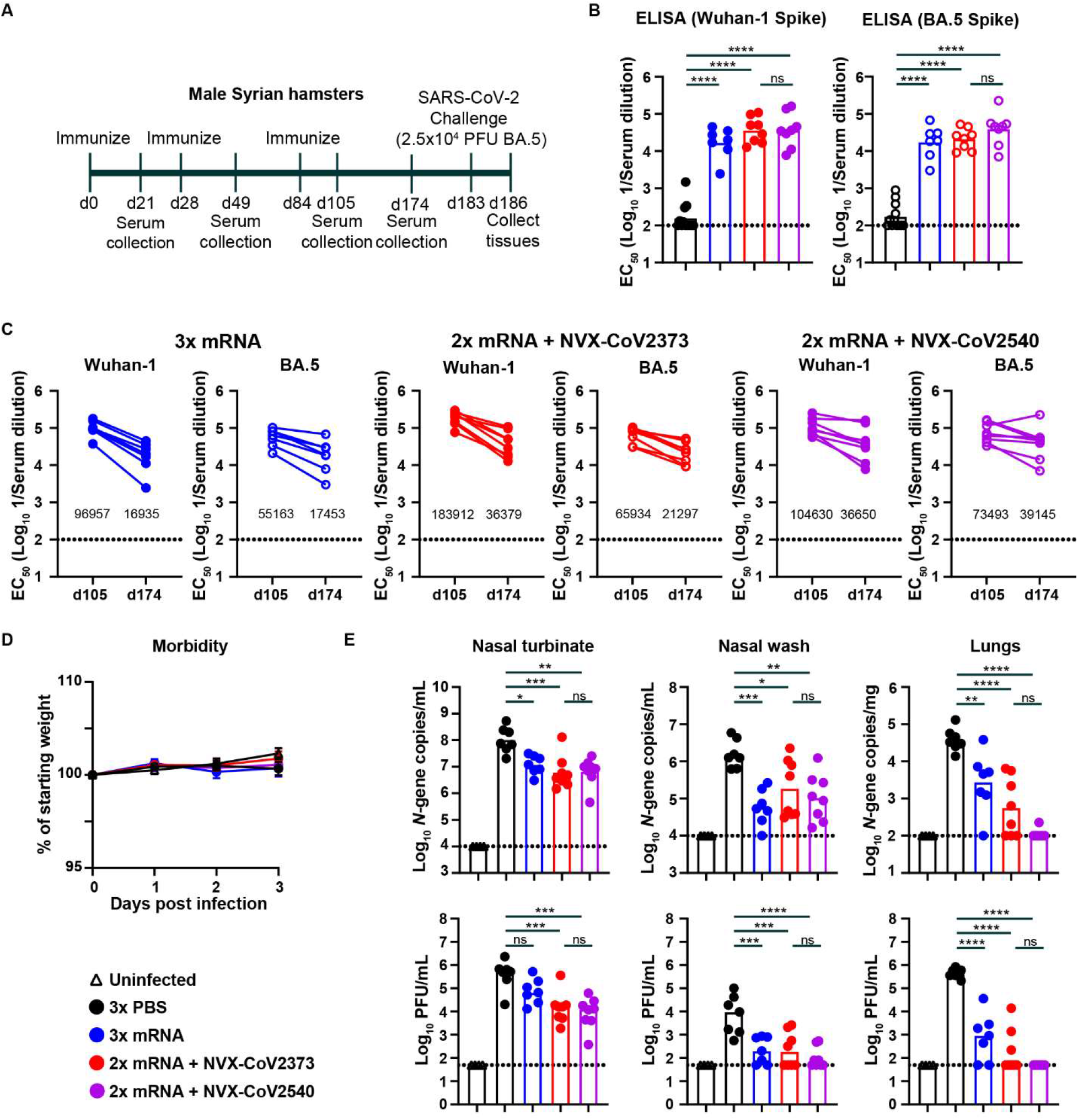
Boosting with NVX-CoV2373 and NVX-CoV2540 provides longer-term protection against challenge with BA.5. (**A**) Experimental design. (**B**) Serum antibody titers (EC_50_) against Wuhan-1 or BA.5 S protein measured by ELISA ∼13 weeks after boosting (third dose) with mRNA (blue symbols), NVX-CoV2373 (red symbols), or NVX-CoV2540 (purple symbols) vaccines. (**** *P* < 0.0001, ns = not significant by one-way ANOVA with a Šídák post-test on ln-transformed EC_50_ values). (**C**) Fold-difference in serum antibody titers (EC_50_) against Wuhan-1 and BA.5 S protein in hamsters immunized twice with mRNA vaccine and boosted with mRNA (blue symbols), NVX-CoV2373 (red symbols), or NVX-CoV2540 (purple symbols) vaccines. Serum was collected ∼13 weeks after the third dose of the vaccine. (**D**) Mean ± SEM of weight loss/gain in SARS-CoV-2 challenged mice. (**E**) Immunized and control Syrian hamsters were challenged with 2.5 x 10^4^ PFU of BA.5 and nasal turbinates, nasal washes, and lungs were collected for analysis of viral RNA levels (top panels) and infectious virus (bottom panels) (**** *P* < 0.0001, *** *P* < 0.001, ** *P* < 0.01, * *P* < 0.05, ns = not significant by one-way ANOVA with Šídák post-test on ln-transformed data). Uninfected control hamsters (open diamonds) were included as negative controls. Bars indicate the geometric mean values, and dotted lines are the limit of detection of the assays. Animals at the limit of detection are arbitrarily assigned this value. The results are from one experiment, and each symbol represents an individual animal (n = 7-8).

Next, the animals that were held for 14 weeks after boosting were challenged with 2.5 × 10^4^ PFU of BA.5. No substantial weight loss was observed in any of the groups of animals, as reported above (**Fig 3D**). In the nasal turbinate of the unvaccinated control group challenged with BA.5, we detected ∼10^8^ copies of the *N* transcript per mL and 4 × 10^5^ PFU/mL (**Fig 3E**). Boosting with mRNA vaccine decreased (∼10-fold, *P* < 0.05) levels of viral RNA and infectious virus (5-fold, *P* = 0.17) in the nasal turbinates (**Fig 3E**). Animals that received boosters of either NVX-CoV2373 or NVX-CoV2540 had significantly less viral RNA (17-fold, *P* < 0.001 and 16-fold, *P* < 0.01 for NVX-CoV2373 and NVX-CoV2540, respectively) and infectious virus (26-fold, *P* < 0.001 and 44-fold, *P* < 0.001 for NVX-CoV2373 and NVX-CoV2540, respectively) in the nasal turbinates. No significant difference in nasal turbinate virus titers or viral RNA was detected between NVX-CoV2373 and NVX-CoV2540 boosted animals.

In the nasal wash of mRNA boosted hamsters, the amounts of viral RNA (27-fold, *P* < 0.001) and infectious virus (33-fold, *P* < 0.001) were significantly lower than in unvaccinated control animals (**Fig 3E**). Similarly, boosting with NVX-CoV2373 or NVX-CoV2540 reduced the levels of viral RNA (8-fold, *P* < 0.05, and 15-fold, *P* < 0.01 for NVX-CoV2373 and NVX-CoV2540, respectively) and infectious virus (32-fold, *P* < 0.001, and 71-fold, *P* < 0.001 for NVX-CoV2373 and NVX-CoV2540, respectively) in the nasal wash after BA.5 virus challenge. We found no significant differences between the NXV-CoV2373 and NVX-CoV2540 boost in the nasal wash.

The amount of infectious virus and *N* gene copy number was also quantified in the lungs. In the unvaccinated control group challenged with BA.5, we detected ∼4 × 10^4^ SARS-CoV-2 *N* gene copies/mg and 4 × 10^5^ PFU/mL (**Fig 3E**). Hamsters that received a booster dose of mRNA vaccine 14 weeks prior to the virus challenge had a lower *N* gene copy number (14-fold, *P* < 0.01) and infectious virus levels (∼550-fold, *P* < 0.0001, **Fig 3E**). Hamsters that received NVX-CoV2373 as the booster had significantly less viral RNA (67-fold, *P* < 0.0001) and infectious virus (∼2,400-fold, *P* < 0.0001), albeit there were more breakthrough infections, defined as positive by RT-qPCR or plaque assay, compared to the BA.5 challenge one month after boosting. Hamsters that were immunized three months earlier with the BA.5 S protein containing vaccine, NVX-CoV2540, had lower viral RNA levels (340-fold, *P* < 0.0001) and infectious viral burden (∼9,000-fold, *P* < 0.0001) in the lungs, with one breakthrough lung infection (**Fig 3E**). The difference in infectious virus and viral RNA levels between NVX-CoV2373 and NVX-CoV2540 boosted animals was not statistically significant. However, the frequency of breakthrough infections in the NVX-CoV2540 boosted animals was significantly lower (*P* < 0.05) compared to NVX-CoV2373 boosted animals three months after vaccination.

## DISCUSSION

In this study, we evaluated the immunogenicity and efficacy of two subunit nanoparticle COVID-19 vaccines, administered as a booster to Syrian hamsters that had received a primary mRNA vaccine series. Heterologous boosting with NVX-CoV2373 or NVX-CoV2540 induced high levels of serum antibodies against ancestral and Omicron BA.5 variant of SARS-CoV-2, respectively. These antibody responses were associated with reduced viral burden after intranasal challenge with the BA.5 variant of SARS-CoV-2. Overall, these data demonstrate the efficacy of these two vaccines as boosters against an Omicron lineage SARS-CoV-2 strain in the pre-clinical hamster model of COVID-19.

Our studies provide immunogenicity and efficacy analysis of NVX-CoV2373 and NVX-CoV2540 in Syrian hamsters. Heterologous boosting with NVX-CoV2373 and NVX-CoV2540 after a primary immunization series with an mRNA vaccine induced significantly higher antibody responses that resulted in less infection than hamsters that received a homologous third booster dose of the mRNA vaccine. Also matched mRNA boosted animals experienced more breakthrough infections in the lungs, as defined by a positive RT-qPCR or plaque assay (5/7 and 6/7 one and three months after receiving the booster dose, respectively). The difference in vaccine efficacy between mRNA and NVX-CoV2373, although not the goal of this study, may be due to differences in antigen dose and vaccine platform. While the mRNA vaccine used in this study are remnant vaccines that were freeze-thawed a second time prior to use, the vaccine was highly immunogenic in hamsters after primary immunization and boosting. We have also shown that the additional freeze-thaw step did not negatively affect the antibody responses to the S protein (**Fig S2**).

A heterologous boost with the NVX-CoV2373 vaccine in hamsters previously immunized with two doses of an mRNA vaccine reduced infection compared to unvaccinated controls. Nonetheless, breakthrough infections in the lungs were detected in hamsters challenged one month (2/7), and three months (5/8), after receiving the third immunization. The frequency of breakthrough infections in the lungs was reduced in hamsters that received the antigen-matched NVX-CoV2540 (BA.5) booster (0/7 at one month and 1/8 three months after the last immunization, respectively; *P* < 0.05, Chi-square test). These data support the use of closely matched or bivalent vaccines to protect against Omicron variants of SARS-CoV-2 ^13,24^. Clearly, additional studies are warranted to assess the durability of boosting effects on breakthrough infections beyond the three-month window that we evaluated. Viral loads were significantly reduced in animals boosted with NVX-CoV-2373 and NVX-CoV2540, however, none of the vaccines protected against infection in the nasal turbinates with nearly 100% of the hamsters having infectious virus present in this tissue. This is consistent with the relatively poor mucosal immune responses generated after administration of intramuscular SARS-CoV-2 vaccines.

### Limitations of the study

We note several limitations of our study. (a) The BA.5 variant of SARS-CoV-2 does not transmit efficiently in hamsters. Accordingly, we did not evaluate the effects of the vaccine on the transmission of SARS-CoV-2, which may be an important measure of vaccine protection. (b) We did not evaluate the efficacy of each vaccine booster in male and female hamsters. Due to the number of variables (vaccines and time after vaccination), testing male and female animals in each experiment was not feasible. (c) Mucosal and T-cell responses after immunization were not measured due to the lack of validated reagents and assay. (d) Studies with more recent emerging variants (*e.g*., XBB.1.5) are warranted.

Overall, our studies demonstrate that boosting of mRNA vaccine responses with the protein nanoparticle vaccines NVX-CoV2373 and NVX-CoV2540 provide protection against a challenge with BA.5 in Syrian hamsters.

## ACKNOWLEDGEMENTS

This study was supported by the NIH (NIAID Center of Excellence for Influenza Research and Response (CEIRR)) contract 75N93021C00016, 75N93021C00014, 75N93019C00051, and R01 AI157155.

## AUTHOR CONTRIBUTIONS

T.L.B. performed hamster experiments. T.L.B. quantified virus titers in collected tissues. A.J. determined viral load by real-time quantitative RT-PCR. N.S. performed ELISA assays. A.C.M.B. analyzed the data. S.M.S. performed virus neutralization assays. A.C.M.B. performed the statistical analysis. A.C.M.B. had unrestricted access to all the data. M.S.D. provided key reagents, supervised experiments, and acquired funding. A.C.M.B. wrote the first draft of the manuscript and all authors reviewed and edited the final version. All authors agreed to submit the manuscript, read and approved the final draft, and take full responsibility of its content.

## SUPPLEMENTARY FIGURES

**Figure S1:**
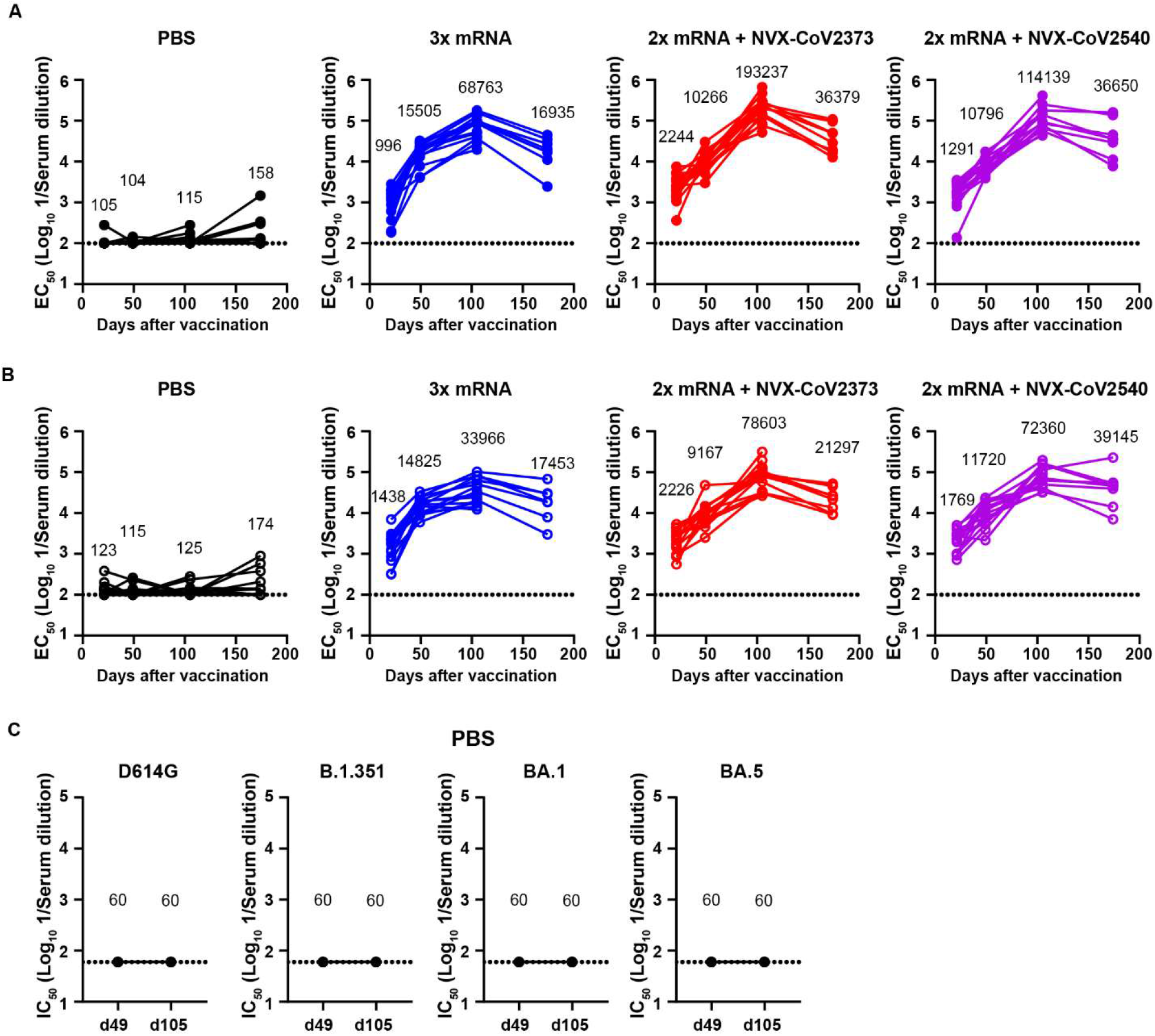
Serum antibody titers against recombinant spike protein. (**A**) Wuhan-1 S protein-specific serum IgG(H+L) titers (EC_50_) in hamsters immunized three times via intramuscular route with PBS (black symbol) or mRNA (blue symbol), or two times with mRNA followed by NVX-CoV2373 (red symbol) or NVX-CoV2540 (purple symbol) boosting. The geometric mean EC_50_ value for each time point is calculated. (**B**) BA.5 S protein-specific serum IgG(H+L) titers in hamsters immunized three times intramuscular with PBS (black symbol) or mRNA (blue symbol), or two times with mRNA followed by NVX-CoV2373 (red symbol) or NVX-CoV2540 (purple symbol) boosting. Each line is an individual animal. (**C**) Serum neutralizing antibody responses (IC_50_) against WA1/2020-D614G, B.1.351, BA.1, or BA.5 SARS-CoV-2 in PBS control hamsters. Serum was collected 21 days after the second and third dose of the vaccine. Dotted line is the limit of detection.

**Figure S2:**
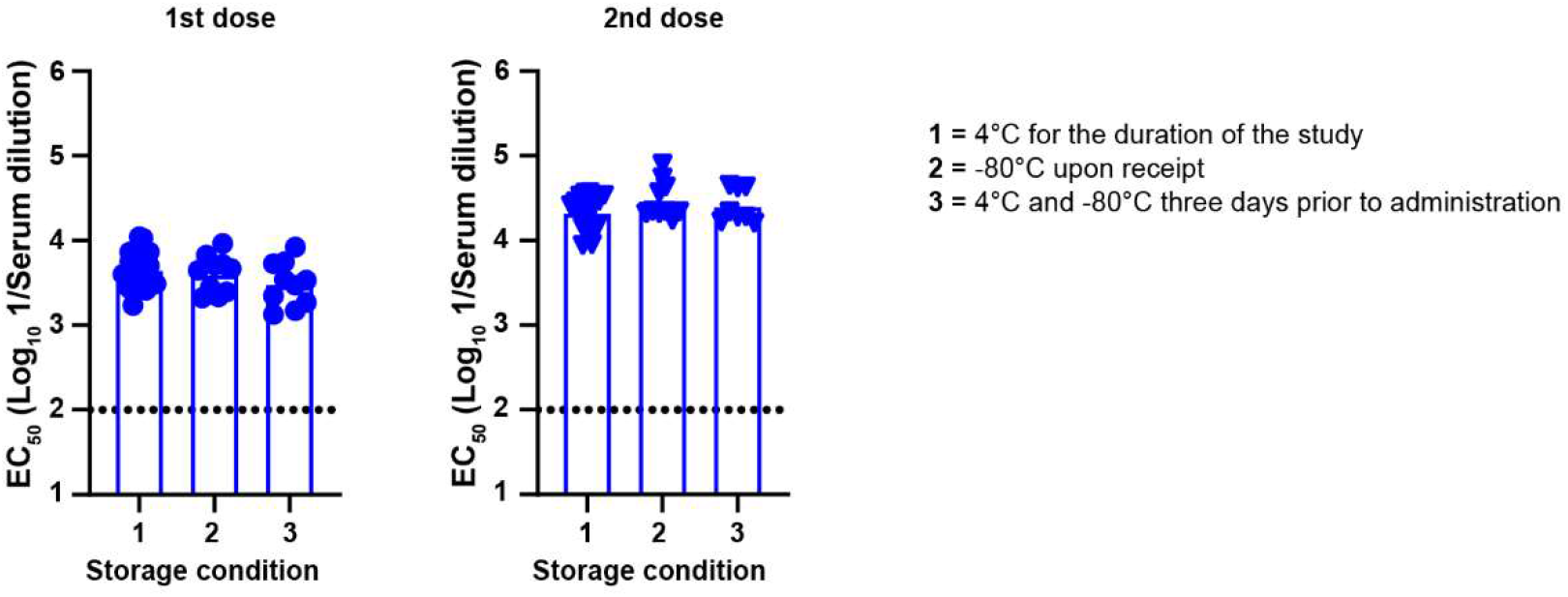
Serum antibody titers against recombinant Wuhan-1 spike protein. Wuhan-1 S protein-specific serum IgG(H+L) titers (EC_50_) in hamsters immunized once (left panel) or twice (right panel) via intramuscular route with remnant 5 µg mRNA vaccine (BNT162b2). The mRNA vaccine was kept at 4°C for the duration of the experiment (#1), stored at -80°C upon receipt and thawed a second time prior to injection (#2), or kept at 4°C and freeze-thawed three days prior to administration (#3). Each dot is an individual animal. Dotted line is the limit of detection.

## STAR METHODS

### RESOURCE AVAILABILITY

#### Lead contact

Further information and requests for resources and reagents should be directed to the Lead Contact, Adrianus C.M. Boon.

#### Materials availability

All requests for resources and reagents should be directed to the Lead Contact author. This includes viruses, vaccines, and primer-probe sets. All reagents will be made available on request after completion of a Materials Transfer Agreement.

#### Data and code availability

All data supporting the findings of this study are available within the paper and are available from the corresponding author upon request. This paper does not include original code. Any additional information required to reanalyze the data reported in this paper is available from the lead contact upon request.

### EXPERIMENTAL MODEL AND SUBJECT DETAILS

#### Cells and Viruses

Vero cells expressing human ACE2 and TMPRSS2 (Vero-hACE2-hTMPRSS2 ^25,26^, gift from Adrian Creanga and Barney Graham, NIH) were cultured at 37°C in Dulbecco’s Modified Eagle medium (DMEM) supplemented with 10% fetal bovine serum (FBS), 10 mM HEPES (pH 7.3), 100 U/mL of Penicillin-Streptomycin, and 10 µg/mL of puromycin. Vero cells expressing TMPRSS2 (Vero-hTMPRSS2) ^26^ were cultured at 37°C in Dulbecco’s Modified Eagle medium (DMEM) supplemented with 10% fetal bovine serum (FBS), 10 mM HEPES (pH 7.3), 100 U/mL of Penicillin-Streptomycin, and 5 µg/mL of blasticidin.

The BA.5 variant of SARS-CoV-2 (hCOV-19/USA/COR-22-063113/2022) was a gift from R. Webby (St. Jude Children’s Research Hospital) and propagated on Vero-hTMPRSS2 cells. The virus stocks were subjected to next-generation sequencing, and the S protein sequences were identical to the original isolates. The infectious virus titer was determined by plaque and focus-forming assay on Vero-hACE2-hTMPRSS2 or Vero-hTMPRSS2 cells.

#### Recombinant proteins

Recombinant S was expressed as previously described ^14,27^. SARS-CoV-2 rS, construct BV2373, is a recombinant nanoparticle vaccine constructed from the full-length, wild-type SARS-CoV-2 spike glycoprotein (GenBank accession number, MN908947; nucleotides 21563–25384). The native full-length S protein was modified by mutation of the putative furin cleavage site RRAR to QQAQ (3Q) located within the S1/S2 cleavage domain to be protease resistant. Two additional proline amino acid substitutions were inserted at positions K986P and V987P (2P) within the heptad repeat 1 (HR1) domain to stabilize SARS-CoV-2 S in a prefusion conformation, which is believed to optimize presentation of neutralizing epitopes. The Omicron BA.5 variant vaccine (construct BV2540) sequence was obtained from the GISAID database (EPI_ISL_12097410.1). To produce construct BV2540, the native full-length S protein was subjected to mutations applied to the ancestral Wuhan-Hu-1 rS plus additional mutations: V3G, T19I, A27S, G142D, V213G, G339D, S371F, S373P, S375F, T376A, D405N, R408S, K417N, N440K, L452R, S477N, T478K, E484A, F486V, Q498R, N501Y, Y505H, D614G, H655Y, N679K, P681H, N764K, D796Y, Q954H, N969K and deletions: ΔL24, ΔP25, ΔP26, ΔH69, ΔV70. The synthetic transgenes were engineered into the baculovirus vector (BV2373 and BV2540) for expression in *Spodoptera frugiperda* (Sf9) insect cells NVX-CoV2373 and NVX-CoV2540 were formulated with Matrix-M adjuvant and stored at 2-8°C.

#### Hamster experiments

Animal studies were carried out in accordance with the recommendations in the Guide for the Care and Use of Laboratory Animals of the National Institutes of Health. The protocols were approved by the Institutional Animal Care and Use Committee at the Washington University School of Medicine (assurance number A3381–01). Five-week old male hamsters were obtained from Charles River Laboratories and housed at Washington University. Five days after arrival, the animals were immunized via intramuscular injection with 5 µg of remnant freeze-thawed mRNA vaccine (BNT162b2) in 100 µL of PBS (50µL per leg). Control animals received PBS alone. Serum samples were obtained 21 days later and one week later the animals were immunized with a second dose of the same vaccine, and serum was collected 21 days later. Approximately one month later (day 84), the animals were randomly divided into three groups and boosted with a third dose of 5 µg mRNA, or 1 µg of the protein nanoparticle vaccine NVX-CoV2373, or NVX-CoV2540, and serum was collected 21 days later. Approximately two days or 10 weeks later, the animals were transferred to the enhanced Biosafety level 3 laboratory and challenged via intranasal route with 2.5 × 10^4^ PFU of Omicron BA.5 variant. Animal weights were measured daily for the duration of the experiment. Three days after challenge, the animals were necropsied, and their lungs and nasal turbinates were collected for virological analysis. These tissues were homogenized in 1 mL of DMEM, clarified by centrifugation (1,000 × *g* for 5 min) and used for viral titer analysis by quantitative RT-PCR (RT-qPCR) using primers and probes targeting the *N* gene, and by plaque assay. A nasal wash was also collected, by inoculating 1 mL of PBS with 0.1% bovine serum albumin into one nostril and collecting the wash from the other nostril. The nasal wash was clarified by centrifugation (2,000 × *g* for 10 min) and used for viral titer analysis by RT-qPCR using primers and probes targeting the *N* gene, and by plaque assay.

### METHOD DETAILS

#### Focus reduction neutralization titer assay (FRNT)

Serial dilutions of serum samples, starting at 1:60, were incubated with 10^2^ focus-forming units (FFU) of different strains of SARS-CoV-2 for 1 h at 37°C. Antibody-virus complexes were added to Vero-hTMPRSS2 cell monolayers in 96-well plates and incubated at 37°C for 1 h. Subsequently, cells were overlaid with 1% (w/v) methylcellulose in Eagle’s Minimal Essential medium (MEM, Thermo Fisher Scientific). Plates were harvested 30 h (WA1/2020 and B.1.351) or 70 h (BA.1 and BA.5) later by removing overlays and fixed with 4% paraformaldehyde (PFA) in PBS for 20 min at room temperature. Plates were washed and sequentially incubated with a pool of anti-S murine antibodies (SARS2–02, −08, −09, −10, −11, −13, −14, −17, −20, −26, −27, −28, −31, −38, −41, −42, −44, −49, −57, −62, −64, −65, −67 and −71 ^28^ and HRP-conjugated goat anti-mouse IgG (Sigma Cat # A8924) in PBS supplemented with 0.1% saponin and 0.1% bovine serum albumin. SARS-CoV-2-infected cell foci were visualized using TrueBlue peroxidase substrate (KPL) and quantitated on an ImmunoSpot microanalyzer (Cellular Technologies).

#### Virus titration assays

Plaque assays were performed on Vero-hACE2-hTRMPSS2 cells in 24-well plates. Lung tissue homogenates or nasal washes were diluted serially by 10-fold, starting at 1:10, in cell infection medium (DMEM + 2% FBS + 100 U/mL of penicillin-streptomycin). Two hundred and fifty microliters of the diluted virus were added to a single well per dilution per sample. After 1 h at 37°C, the inoculum was aspirated, the cells were washed with PBS, and a 1% methylcellulose overlay in MEM supplemented with 2% FBS was added. Ninety-six hours after virus inoculation, the cells were fixed with 4% formalin, and the monolayer was stained with crystal violet (0.5% w/v in 25% methanol in water) for 30 min at 20°C. The number of plaques were counted and used to calculate the plaque forming units/mL (PFU/mL).

To quantify viral load in lung tissue homogenates and nasal washes, RNA was extracted from 100 µL samples using MagNA Pure 24 Total NA Isolation Kit on the MagNA Pure 24 system (Roche) and eluted with 50 µL of water. Four microliters RNA was used for real-time RT-qPCR to detect and quantify *N* gene of SARS-CoV-2 using TaqMan™ RNA-to-CT 1-Step Kit (Thermo Fisher Scientific) as described ^29^ using the following primers and probes: Forward: GACCCCAAAATCAGCGAAAT; Reverse: TCTGGTTACTGCCAGTTGAATCTG; Probe: ACCCCGCATTACGTTTGGTGGACC; 5’Dye/3’Quencher: 6-FAM/ZEN/IBFQ. Viral RNA was expressed as *N* gene copy numbers per mg for lung tissue homogenates or mL for nasal swabs and nasal washes, based on a standard included in the assay, which was created via *in vitro* transcription of a synthetic DNA molecule containing the target region of the *N* gene.

#### ELISA

Ninety-six-well microtiter plates (Nunc MaxiSorp; ThermoFisher Scientific) were coated with 100 µL of recombinant SARS-CoV-2 S protein (Wuhan-1 strain and BA.5, generated by Novavax as described above) at a concentration of 1 µg/mL in PBS (Gibco) at 4 °C overnight; negative control wells were coated with 1 µg/mL of BSA (Sigma). Plates were blocked for 1.5 h at room temperature with 280 µL of blocking solution (PBS supplemented with 0.05% Tween-20 (Sigma) and 10% FBS (Corning)). The sera were diluted serially in blocking solution, starting at 1:100 dilution and incubated for 1.5 h at room temperature. The plates were washed three times with T-PBS (1X PBS supplemented with 0.05% Tween-20), and 50 µL of HRP-conjugated anti-hamster IgG(H+L) antibody (Southern Biotech Cat. #6061-05) diluted 1:1000 in blocking solution, was added to all wells and incubated for 1 h at room temperature. Plates were washed 3 times with T-PBS and 3 times with 1X PBS, and 50 µL of 1-step Ultra TMB-ELISA substrate solution (Thermo Fisher Scientific) was added to all wells. The reaction was stopped after 10 min using 50 µL of 1N H_2_SO_4_, and the plates were analyzed at a wavelength of 450 nm using a microtiter plate reader (BioTek).

### QUANTIFICATION AND STATISTICAL ANALYSES

Statistical significance was assigned when *P* values were < 0.05 using GraphPad Prism version 9.3. Tests, number of animals, median and geometric mean values, and statistical comparison groups are indicated in the Figure legends. Analysis of weight change was determined by two-way ANOVA. Changes in infectious virus titer, viral RNA levels, or serum antibody responses were compared to unvaccinated immunized animals, and between the two NVX vaccines, and were analyzed by one-way ANOVA with multiple comparisons correction on ln-transformed data.

